# Choosing and learning: outcome valence differentially affects learning from free versus forced choices

**DOI:** 10.1101/637157

**Authors:** Valérian Chambon, Héloïse Théro, Marie Vidal, Henri Vandendriessche, Patrick Haggard, Stefano Palminteri

## Abstract

Positivity bias refers to learning more from positive than negative events. This learning asymmetry could either reflect a preference for positive events in general, or be the upshot of a more general, and perhaps, ubiquitous, “choice-confirmation” bias, whereby agents preferentially integrate information that confirms their previous decision. We systematically compared these two theories with 3 experiments mixing free- and forced-choice conditions, featuring factual and counterfactual learning and varying action requirements across “go” and “no-go” trials. Computational analyses of learning rates showed clear and robust evidence in favour of the “choice-confirmation” theory: participants amplified positive prediction errors in free-choice conditions while being valence-neutral on forced-choice conditions. We suggest that a choice-confirmation bias is adaptive to the extent that it reinforces actions that are most likely to meet an individual’s needs, i.e. freely *chosen* actions. In contrast, outcomes from *unchosen* actions are more likely to be treated impartially, i.e. to be assigned no special value in self-determined decisions.

## Introduction

Standard reinforcement learning models conceive agents as impartial learners: they learn equally well from positive and negative outcomes alike (Barto and Sutton, 1998). However, empirical studies have recently come to challenge this view by demonstrating that human learners, rather than processing information impartially, consistently display a *valence-induced* bias: when faced with uncertain choice options, they tend to disregard bad news by integrating worse-than-expected outcomes (positive prediction errors) at a lower rate relative to better-than-expected ones (negative prediction errors) (Lefebvre et al. 2017; Aberg et al., 2016; Frank et al., 2007). Importantly, this “positivity” bias would echo the asymmetric processing of self-relevant information in probabilistic reasoning, whereby good news on average receives more weight than bad news (Sharot and Garrett, 2016; Kuzmanovic et al., 2018).

A bias for learning preferentially from better-than-expected outcomes would reflect a preference for positive events *in general*. This prediction is however at odds with recent findings. In a two-armed bandit task featuring *complete* feedback information, we previously found that participants would learn preferentially from better-than-expected *obtained* outcomes, while preferentially learning from worse-than-expected *forgone* outcomes, i.e., from the outcome associated with the option they had not chosen (Palminteri et al., 2017).

This learning asymmetry suggests that what has been previously characterized as a “positivity” bias may, in fact, be the upshot of a more general, and perhaps ubiquitous, “choice-confirmation” bias, whereby human agents preferentially integrate information that confirms their previous decision (Nickerson, 1998).

Building on these previous findings, we reasoned that if human reinforcement learning is indeed biased in a ‘choice-confirmatory’ manner, learning from action-outcome couplings that were not *voluntarily chosen* by the subject (“forced choice”) should present no bias. To test this hypothesis, we conducted two experiments involving instrumental learning and computational model-based analyses. Participants were administrated new variants of a probabilistic learning task in which they could either “freely” choose between two options, or were “forced” to implement the choice made by a computer. In the first experiment (N = 24), participants were only shown the obtained outcome corresponding to their choice (factual learning). In the second experiment (N = 24), participants were shown both the obtained and the forgone outcome (counterfactual learning). We had two key predictions. With regard to *factual* learning, participants should learn better from positive prediction error, but they should only do so when *free* to choose (free-choice trials), while showing no effect when forced to match a computer’s choice (forced-choice trials). With regard to *counterfactual* learning from forgone outcomes, we expect the opposite pattern: in free-choice trials, negative prediction errors should be more likely to be taken into account than positive prediction errors, while we expect no bias in forced-choice trials. Put another way, we expect to observe a confirmation bias only when outcomes derive from self-determined choices.

To verify our predictions, we fitted subjects’ behavioural data with several variants of reinforcement learning model, including different learning rates as a function of whether the outcome was positive or negative, obtained or forgone, and followed a free or a forced choice. Learning rate analyses were coupled with model comparison analyses aimed at evaluating evidence for the current hypothesis (i.e., no confirmation bias in observational learning) and ruling out alternative interpretations of the results (i.e., a perseveration bias, see Katahira, 2018).

Another central question regarding the nature of the valence-induced bias concerns its relation with action requirements. An influential theory and related previous findings suggest that positive outcomes favour learning of choices involving action execution, while negative outcomes would favour learning of choices involving action *withdrawal* (e.g., Boureau and Dayan, 2011; Guitart-Masip et al., 2012). Extending this framework to feedback processing, one should expect the positivity bias to disappear, or even reverse, following trials in which decision is made by “refraining” from action. However, if the positivity bias emerges as a consequence of choice confirmation, then only making a choice (vs. following an instruction) should matter, irrespective of whether this choice is executed through making an action or not. Using a modified version of our design, we tested this prediction in a third experiment that varied the requirements of motor execution by including both “go” and “no-go” trials (N=24). Learning rates were analysed as function of both outcome valence (negative vs. positive) and the requirement for motor execution in order to implement the selected action (key press vs. no key press).

## Results

Participants performed instrumental learning tasks, involving free- and forced-choice trials, and “go” or “no-go” trials (see **Methods** and **Table 1**). The task consisted in cumulating as many points as possible by selecting whichever of two symbols was associated with the highest probability of reward. Symbols were always presented in pairs, which comprised one more rewarding and one less rewarding option. In all experiments, each block was associated with a specific pair of symbols, meaning that the participant had to learn from scratch the reward contingencies at the beginning of each block.

**Table 1.**
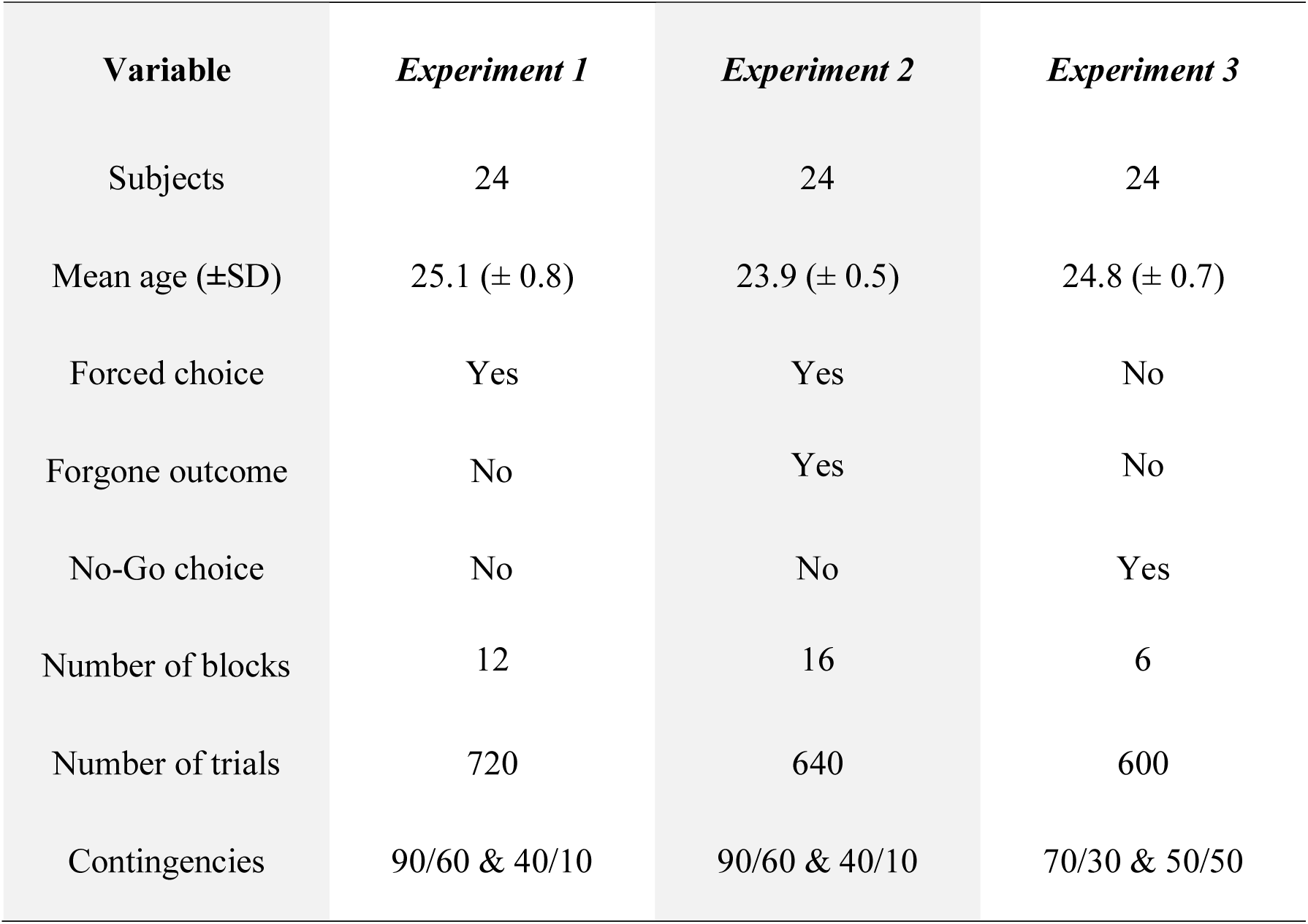
Sociodemographic and design information for the three experiments.

| **Variable** | ***Experiment 1*** | ***Experiment 2*** | ***Experiment 3*** |
| --- | --- | --- | --- |
| Subjects | 24 | 24 | 24 |
| Mean age (±SD) | 25.1 (± 0.8) | 23.9 (± 0.5) | 24.8 (± 0.7) |
| Forced choice | Yes | Yes | No |
| Forgone outcome | No | Yes | No |
| No-Go choice | No | No | Yes |
| Number of blocks | 12 | 16 | 6 |
| Number of trials | 720 | 640 | 600 |
| Contingencies | 90/60 & 40/10 | 90/60 & 40/10 | 70/30 & 50/50 |

In the first two experiments, free-choice trials were interleaved with forced-choice trials. In the latter, the computer randomly preselected a symbol, forcing the participant to match the computer’s choice (**Figure 1A**). Experiment 1 (N = 24) featured “partial” feedback information, since only the obtained outcome (i.e., the outcome of the chosen symbol) was shown (“Exp.1”, **Figure 1A**, top panel). Experiment 2 (N = 24), featured “complete” feedback information, since both the obtained and forgone outcomes (i.e., the outcome of the unchosen symbols) were shown (“Exp.2”, **Figure 1A**, bottom panel). In Experiment 3 (N = 24), action requirements were varied within trials where the choice could either be made by performing an action (key press, “go” trials) or by refraining from acting (no key press, “no-go” trials) (“Exp.3”, **Figure 1B**).

**Figure 1.**
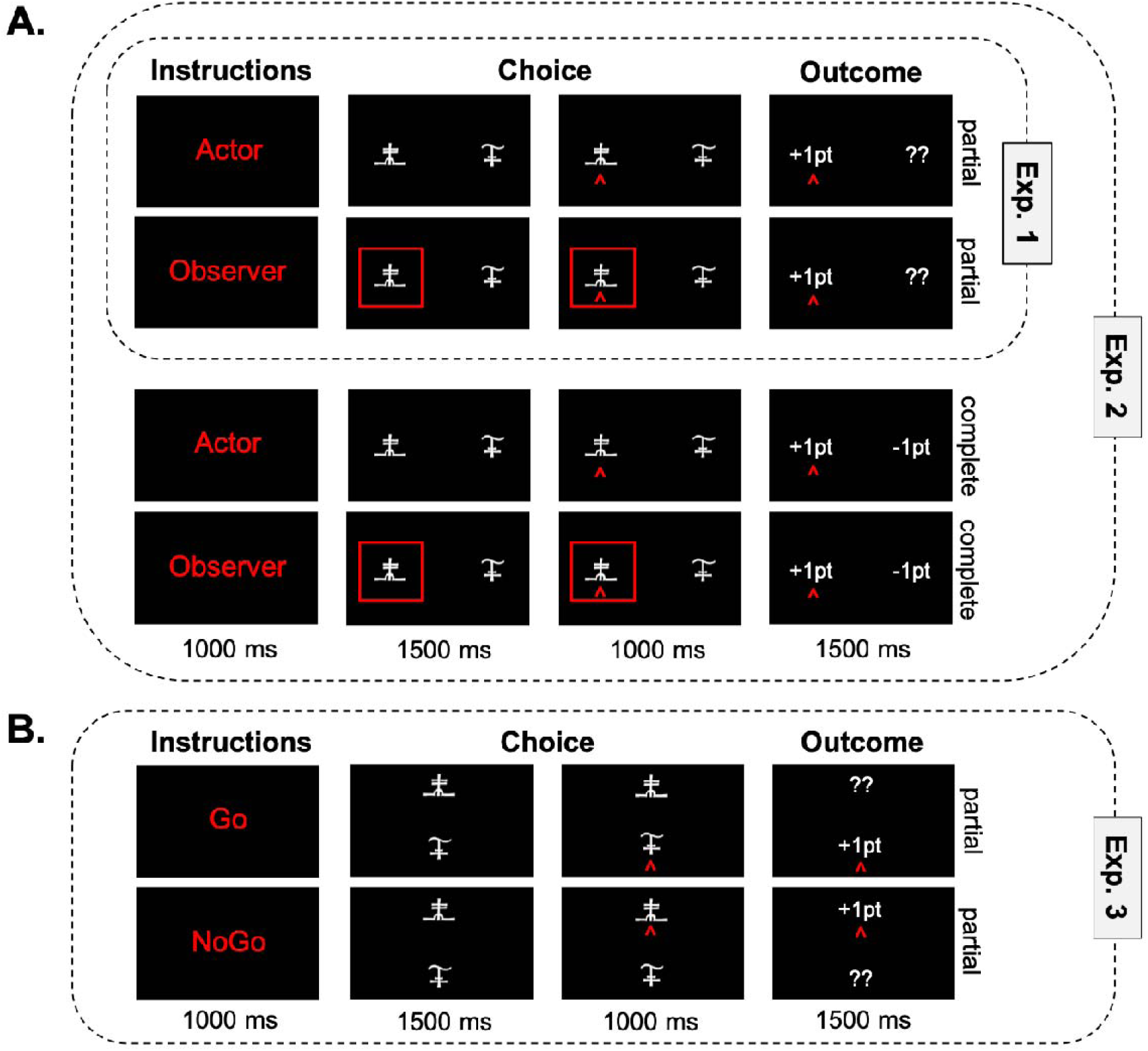
Schematic of trial procedure and stimuli. **(A)** Description of the four trial types implemented in Experiments 1 (top panel) and Experiment 2 (bottom panel). In free-choice trials (“Actor”), participants could freely choose between two options, while in forced-choice trials (“Observer”) participants had to match a preselected option, which was indicated by a red square. In “partial” trials, participants were only shown the outcome (+1 or -1) associated to the chosen option, while in “complete” trials participants were shown the outcomes associated to both chosen and unchosen options. Experiment 1 included a condition with only free-choice trials, and a condition with intermixed free- and forced-choice trials. Only “partial” trials were used. In Experiment 2, free- and forced-choice trials were intermixed, within two conditions: one with “partial” trials, and one with “complete” trials, where the outcomes of both chosen and unchosen options were shown. **(B)** Description of the two conditions implemented in Experiment 3. Action requirements were varied within trials where the choice of an option could either be made by pressing a key (“Go” trials) or by refraining from pressing any key (“No-Go” trials). This experiment only featured “free-choice” trials and “partial” feedback.

## Learning performance

To verify that participants understood the task correctly, we analysed correct choice rate (i.e., the rate of choosing the most rewarding symbol) and found it significantly higher than chance level in all 3 experiments (two-tailed t-tests against 50%: all *t’s* > 10, all *p’s* < 10^-8^). To assess learning dynamics, we also verified that learning performance was higher in the second, relative to the first, half of the learning block (see **Figure 2A**).

**Figure 2.**
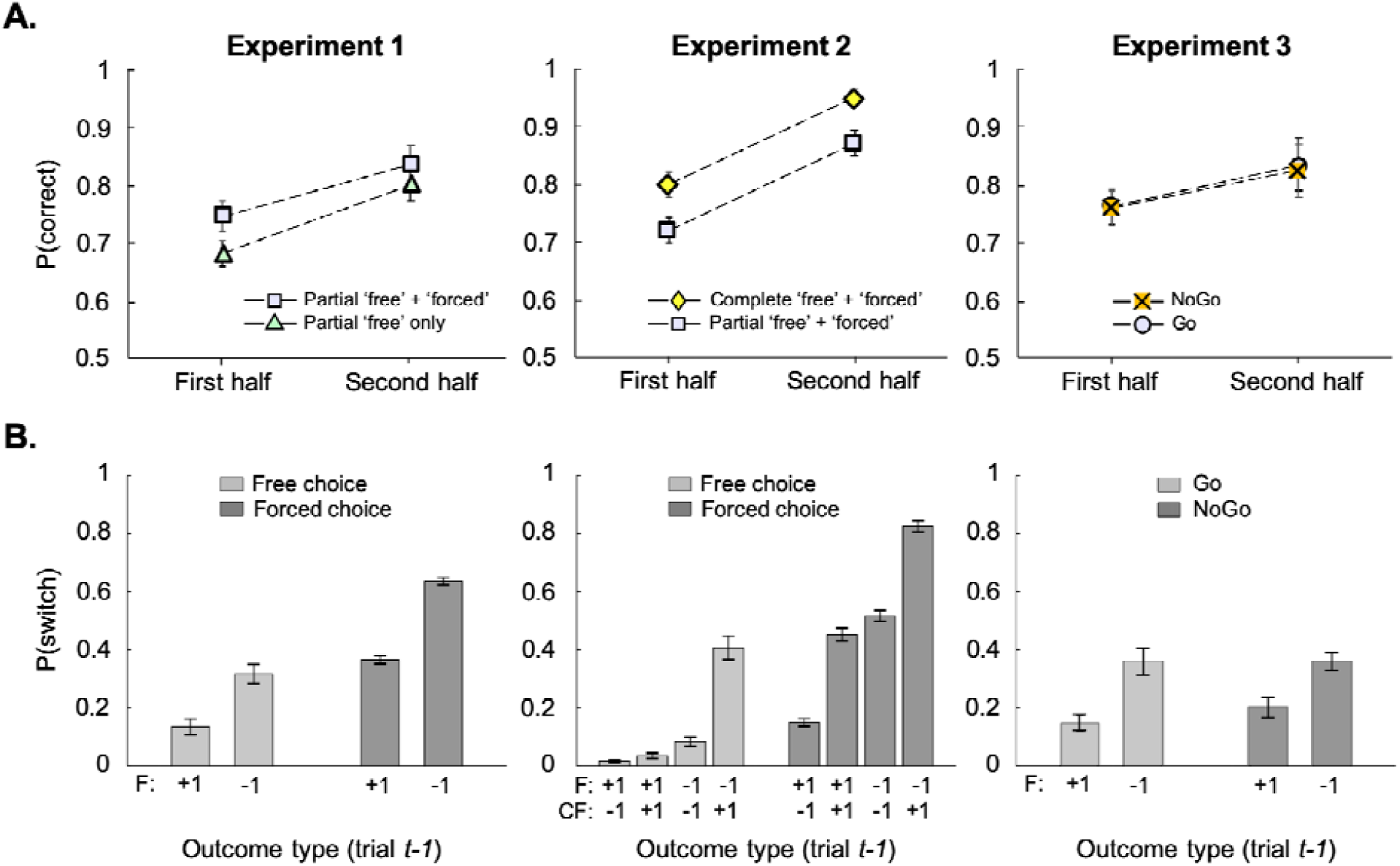
Behavioural results. **(A)** Mean proportion of correct choices in the first and second halves of each learning block, for the two conditions (free only; intermixed free and forced) of Experiment 1 (left) and for the two conditions (partial; complete) of Experiment 2 (middle), and for go and no-go trials in Experiment 3 (right). In Experiment 3, the proportion of correct choices could only be analysed for the 70/30 (vs. 50/50) contingencies. **(B)** Proportion of choice switches between trial *t* and *t*-1 as a function of the obtained outcome (factual, *F*) and the forgone (counterfactual, *CF*) outcome seen on trial *t*-1, depending on whether this trial was a free- or a forced-choice trial (Exp. 1 and 2), or a go or a no-go trial (Exp. 3). For the Experiment 1, the analysis was made on “partial” intermixed trials. For the Experiment 2, the analysis was made on “complete” intermixed trials, which contained both obtained and forgone outcomes. Error bars are standard errors calculated across participants.

Switching choice after a negative outcome, and repeating a choice after receiving a positive outcome, is a hallmark of feedback-based adaptive behaviour. To verify that participants took into account both free and forced choice outcomes, we analysed the switch rate as a function of switches depending on (i) whether the previous obtained outcome was positive or negative, but also on (ii) whether the previous trial was a free- or forced-choice trial (Exp. 1 and 2) or a go or a no-go trial (Exp. 3), and on (iii) whether the *forgone* outcome was positive or negative (Experiment 2 only).

The repeated-measures ANOVAs revealed a main effect of the *obtained* outcome on switch choices in all experiments (all *F’s* > 16, all *p’s* < 10^-3^). Thus, as expected, participants switched more often options after receiving a negative, relative to a positive, outcome. This effect was observed after both a free- and a forced-choice trial alike, and in both go and no-go trials. As expected, we also found a main effect of the *forgone* outcome in Experiment 2 (*F_1,23_* = 39, *p* = 3.6 × 10^-9^), with participants switching choices significantly more when the outcome associated with the *unchosen* option was *positive*, relative to negative (see **Figure 2B**). Finally, in both Experiments 1 and 2 the main effect of the type of choice was significant (all *F’s* > 36, all *p’s* < 10^-7^), with participants switching more often after a forced-choice trial than after a free-choice trial. This effect can be accounted for by the fact that the chosen symbol was pseudo-randomly selected in forced-choice trials, while subjects preferentially chose the “correct” option in free-choice trials – thus forced-choice trials were more likely to involve incorrect choices. In Experiment 3, neither the main effect of the execution mode (key press *vs.* no key press) or the execution-by-valence interaction effect was significant (all *p’s* > 0.05).

## Model parameter analyses

To test the influence of outcome valence and choice type on learning, we fitted the data with a modified Rescorla-Wagner model assuming different learning rates for (i) positive and negative outcomes (α^+^ and α^-^) and (ii) for free- and forced-choice trials (Experiments 1 & 2) or (ii) for Go and No-Go trials (Experiment 3). In Experiment 2, different learning rates were also assumed for (iii) obtained and forgone outcomes (**Figure 3**, bottom). We refer to these models as “full” models, because they present the highest number of parameters in the considered model space. In Experiment 1, the resulting learning rates were subjected to a 2 × 2 repeated-measures ANOVA with outcome valence (positive vs. negative) and choice type (free vs. forced) as within-subject factors. In Experiment 2, learning rates were subjected to a 2 × 2 × 2 repeated-measures ANOVA with outcome valence (positive vs. negative), choice type (free vs. forced), and outcome type (obtained vs. forgone), as within-subject factors. Finally, in Experiment 3, learning rates were subjected to a 2 × 2 repeated-measures ANOVA with outcome valence (positive vs. negative) and execution mode (key press vs. no key press), as within-subject factors.

**Figure 3.**
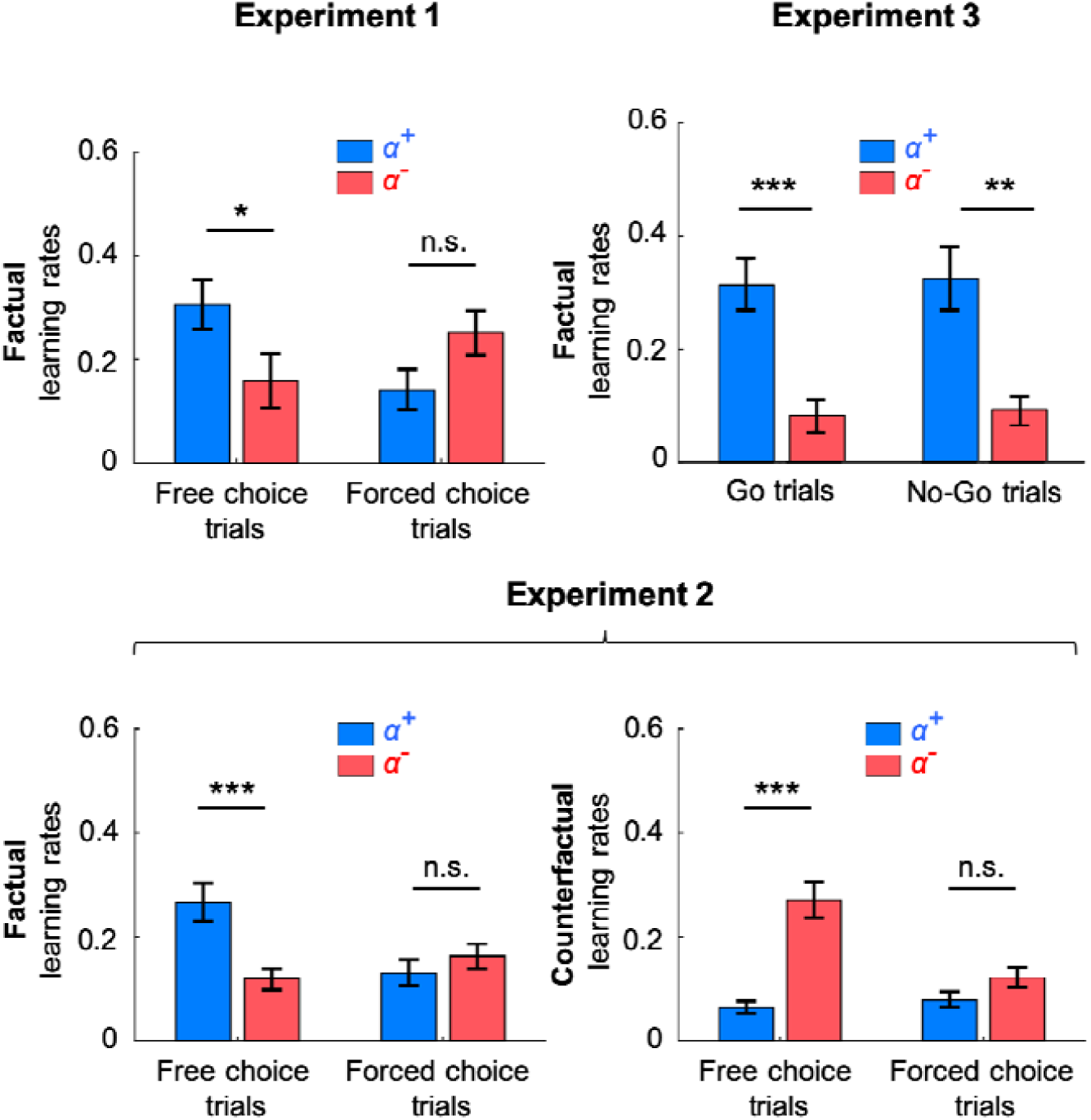
Parameter results of the “full” model from all 3 experiments. Top left panel: Fitted *factual* learning rates from free- and forced-choice trials of Experiment 1. *Bottom panel:* Fitted *factual* and *counterfactual* learning rates from free- and forced-choice trials of Experiment 2. *Top right panel:* Fitted learning rates from Go and No-Go trials of Experiment 3. Note that only obtained outcomes were shown in Experiments 1 and 3, whereas both obtained and forgone outcomes were displayed in Experiment 2, which allowed for fitting counterfactual learning rates. Positive (α^+^) and negative (α^-^) learning rates were represented in blue and red, respectively. N.s.: *p* > 0.05; *: *p* < 0.05; **: *p* < 0.01; ***: *p* < 0.001.

In both Experiments 1 and 2, no main effect was significant. In Experiment 3, only the main effect of outcome valence was significant (*F_1,23_* = 26,8, *p <* 1.0 × 10^-3^). In Experiment 1, we found a significant valence-by-choice interaction (*F_1,23_* = 7.4, *p* = 7.6 × 10^-3^). We found a significant valence-by-outcome (*F_1,23_*= 11, p = 1.2 × 10^-3^) and a significant valence-by-choice-by-outcome (*F_1,23_* = 6.8, *p* = 1.0 × 10^-2^) interaction in Experiment 2. The valence-by-execution-mode interaction was not significant in Experiment 3 (**Figure 3** and **Table 2**). We performed post-hoc t-tests to further investigate significant interactions found in Experiments 1 and 2. The difference between positive and negative learning rates was significant in free-choice trials for both experiments (Exp. 1, obtained outcomes: *t_23_* = 2.5, *p* = 2.0 × 10^-2^; Exp. 2, obtained outcomes: *t_23_* = 4.1, *p* = 4.3 × 10^-4^; forgone outcomes: *t_23_* = -6.2, p = 2.6 × 10^-6^), but was not significant in forced-choice trials (Exp. 1, obtained: *t_23_*= -2.0, *p* = 0.055; Exp. 2, obtained: *t_23_* = -1.3, *p* = 0.20; and forgone: *t_23_* = -1.5, *p* = 0.14, see **Figure 3**).

**Table 2.** Mean parameter values (± standard errors) of the winning model in all 3 experiments.

|  | *Param.* | *Factual* |  |  | *Counterfactual* |  |  |
| --- | --- | --- | --- | --- | --- | --- | --- |
| | $\beta$ | $\alpha^+_{\text{free}}$ | $\alpha^-_{\text{free}}$ | $\alpha_{\text{forced}}$ | $\alpha^+_{\text{free}}$ | $\alpha^-_{\text{free}}$ | $\alpha_{\text{forced}}$ |
| **Exp. 1** | 4.2 | 0.35 | 0.14 | 0.13 | - | - | - |
| | ($\pm 0.50$) | ($\pm 0.063$) | ($\pm 0.054$) | ($\pm 0.036$) | | | |
| **Exp. 2** | 6.3 | 0.30 | 0.11 | 0.14 | 0.065 | 0.27 | 0.089 |
| | ($\pm 0.60$) | ($\pm 0.035$) | ($\pm 0.020$) | ($\pm 0.022$) | ($\pm 0.011$) | ($\pm 0.033$) | ($\pm 0.011$) |
|  |  | *Go trials* |  | *NoGo trials* |  |  |  |
| | $\beta$ | $\alpha^+_{\text{Go}}$ | $\alpha^-_{\text{Go}}$ | $\alpha^+_{\text{NoGo}}$ | $\alpha^-_{\text{NoGo}}$ | | |
| **Exp. 3** | 4.0 | 0.34 | 0.09 | 0.35 | 0.10 | - | - |
| | ($\pm 0.97$) | ($\pm 0.05$) | ($\pm 0.03$) | ($\pm 0.06$) | ($\pm 0.03$) | | |

To sum up, we replicate that participants learned preferentially from positive compared, to negative, prediction errors, whereas the opposite was true for forgone outcomes (Palminteri et al., 2017). Crucially, we found that this learning asymmetry was significant only in free-choice trials, and was undetectable when participants were forced to match the computer’s decision. Varying action requirements across go and no-go trials had no effect on learning asymmetry.

## Parsimony-driven parameter reduction

Although we found no valence-induced bias in forced-choice learning rates on average, one cannot rule out that participants had opposite – significant – biases (e.g., some would learn better from positive forced-choice outcomes, while some others would learn better from negative forced-choice outcomes). We therefore ran a parsimony-driven parameter reduction to assess whether fitting different learning rates in (i) forced-choice trials and (ii) go and no-go trials, better predicted participants’ data (see **Figure 4A**). The full models (i.e., the model with 4 learning rates (α) in experiments 1 and 3, and 8 learning rates (α) in experiment 2) were compared with reduced versions including either a valence-induced bias only for free-choice outcomes or no bias at all. We compared the models using a Bayesian model selection procedure (Daunizeau et al., 2014) based on the Bayesian Information Criterion (BIC). In all experiments, “intermediate” models (i.e., models including valence-induced bias only for free-choice outcomes) were found to better account for the data: their average posterior probabilities were higher than the posterior probabilities of the other models in the set (see **Figure 4B**).

**Figure 4.**
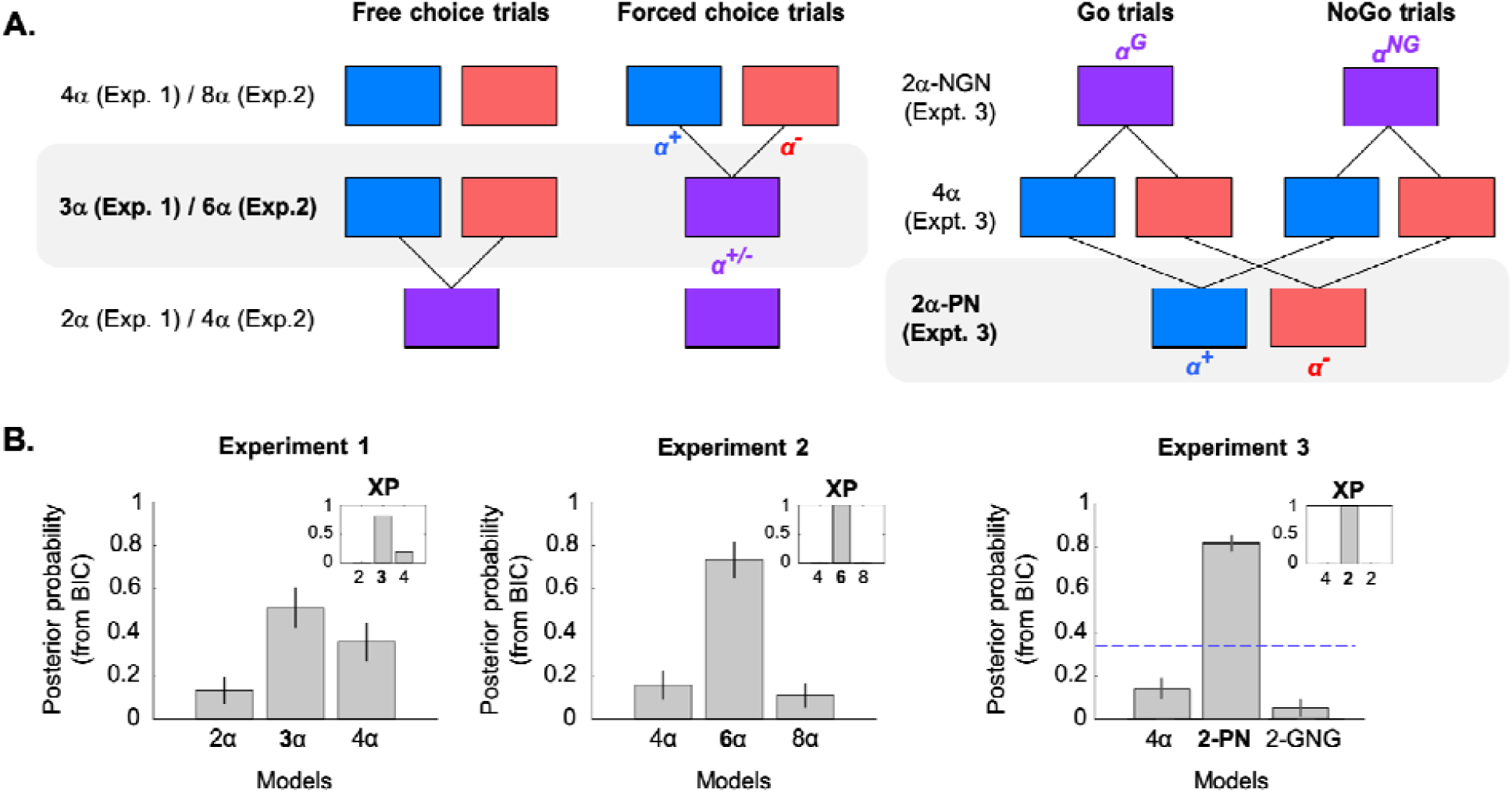
Model comparison results from all 3 experiments. **(A)** Representation of the model space. *Top left and middle panels:* In experiments 1 and 2, the “full” model (4*α* and 8*α*, respectively) has different learning rates for positive and negative prediction errors (blue and red squares, respectively), and for free- and forced-choice trials. The “intermediate” model (3*α* and 6*α*, grey rectangle) has only single learning rate on forced-choice trials, whereas the “reduced” model (2*α* and 4*α*) does not split learning rates by valence at all. The “reduced” model is thus nested within the “intermediate” model, which is itself nested within the “full” model. Note that in Experiment 2, the parameter reduction operates for both factual and counterfactual learning rates. The “intermediate” model is the winning model. *Top right panel:* In experiment 3, the “full” model (4*α*) has different learning rates for positive and negative prediction errors (blue and red squares), and for Go and No-Go trials. Two reduced models were tested: (i) a model with 2 different learning rates for positive and negative outcomes (2*α*-PN), and (ii) a model with 2 different learning rates for Go and No-Go trials (2*α*-GNG). ‘PN’: Positive-Negative; ‘GNG’: Go-NoGo. **(B)** The expectations and the variance of the posterior probability for each model, based on the Bayesian Information Criterion (BIC) values, with the exceedance probability (XP) for each model shown in small inserts. The winning model is indicated in bold.

Consistent with a previous study (Palminteri et al. 2017), we further found that Experiment 2 data were better explained by an even more parsimonious model assuming similar *positive factual* and *negative counterfactual* learning rates 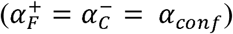, and similar *negative factual* and *positive counterfactual* learning rates 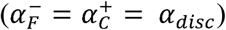. This final model thus had 3 different learning rates: *α_conf_*, *α_disc_*, *α_forced_* (**Figure 5**). We refer to these learning rates as *α_conf_* and *α_disc_* because they embody learning from *confirmatory* (positive obtained and negative forgone) and *disconfirmatory* (negative obtained and positive forgone) outcomes, respectively.

**Figure 5.**
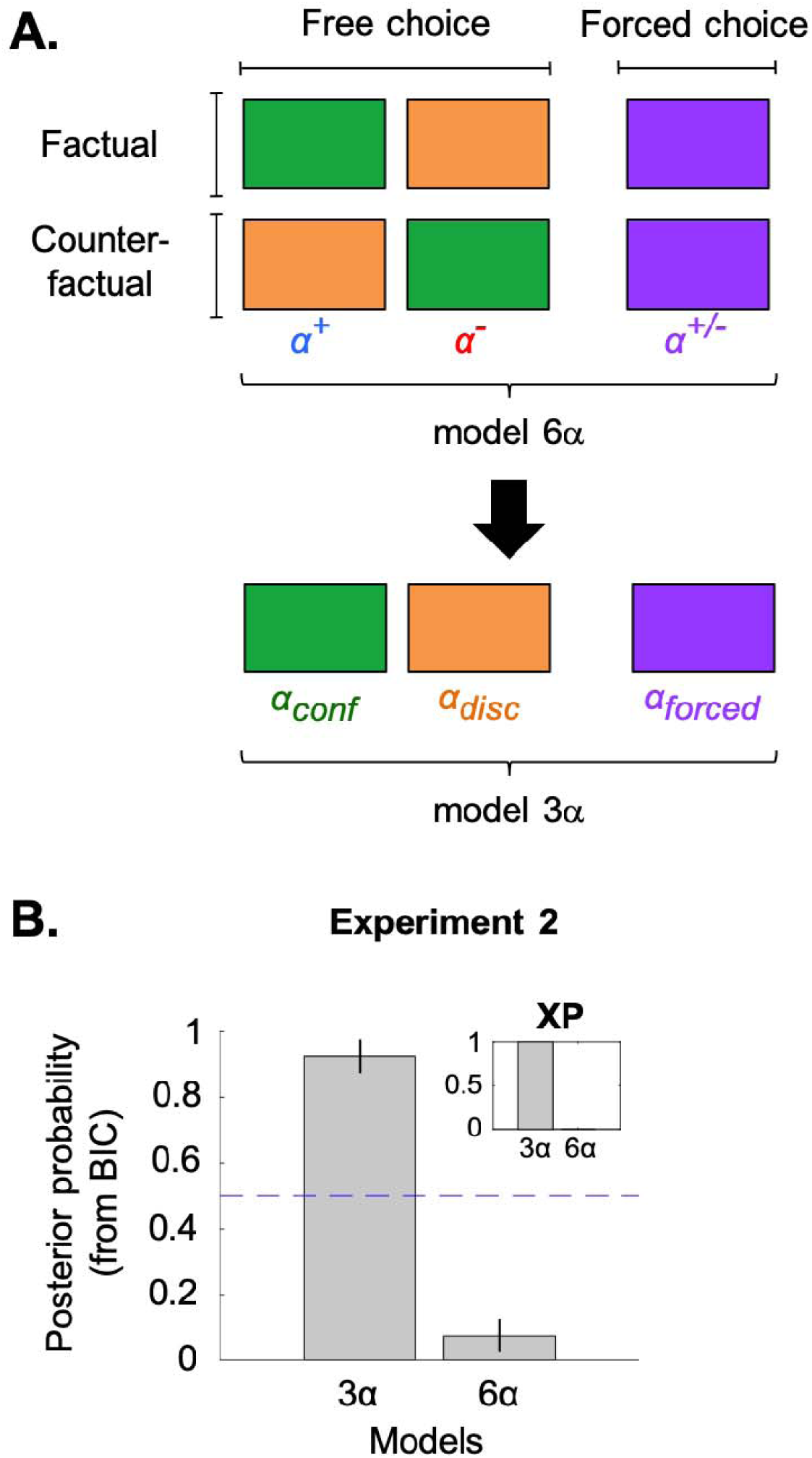
Model comparison from Experiment 2. **(A)** Representation of the model space. The model with 6 learning rates (top) corresponds to the winning model presented in Figure 4A (left panel). The black arrow indicates that this winning model can be reduced to a model with 3 learning rates, which embody learning in forced-choice trials (*_forced_*) and learning from confirmatory (*_conf_*) and disconfirmatory (*_disc_*) outcomes in free-choice trials. **(B)** Expectation and variance of the posterior probability for the initial winning model (6) and the “reduced” model (3), based on the Bayesian Information Criterion (BIC) values. The insert chart shows the exceedance probability (XP) for each model.

## Comparison with the perseveration model

In previous studies, a heightened choice hysteresis (i.e., an increased tendency to repeat a choice above and beyond outcome-related evidence) has been identified as a behavioral hallmark of positivity and confirmatory learning biases (Lefebvre et al., 2017; Palminteri et al., 2017). However, the same behavioral phenomenon may arise in the presence of a learning-independent choice repetition bias (often referred to as “perseveration” bias; see Correa et al., 2018), which is not to confound with motor inertia (which is avoided in our tasks by counterbalancing the spatial position of the cues). Even more concerning, positivity and confirmation biases may spuriously arise fitting multiple learning rates on data presenting a simple choice repetition bias (Katahira, 2018). To rule out this possibility we explicitly compared models including *positivity* 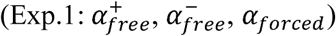 and *confirmation* (Exp.2: *α_conf_*, *α_disc_*, *α_forced_*) learning biases to a model including a *perseveration* parameter. The models with different learning rates were found to better account for the data compared to the perseveration model, with a higher average posterior probability (exceedance probabilities: 0.87 and 1.00 in Experiment 1 and Experiment 2, respectively) (**Figure 6A**).

**Figure 6.**
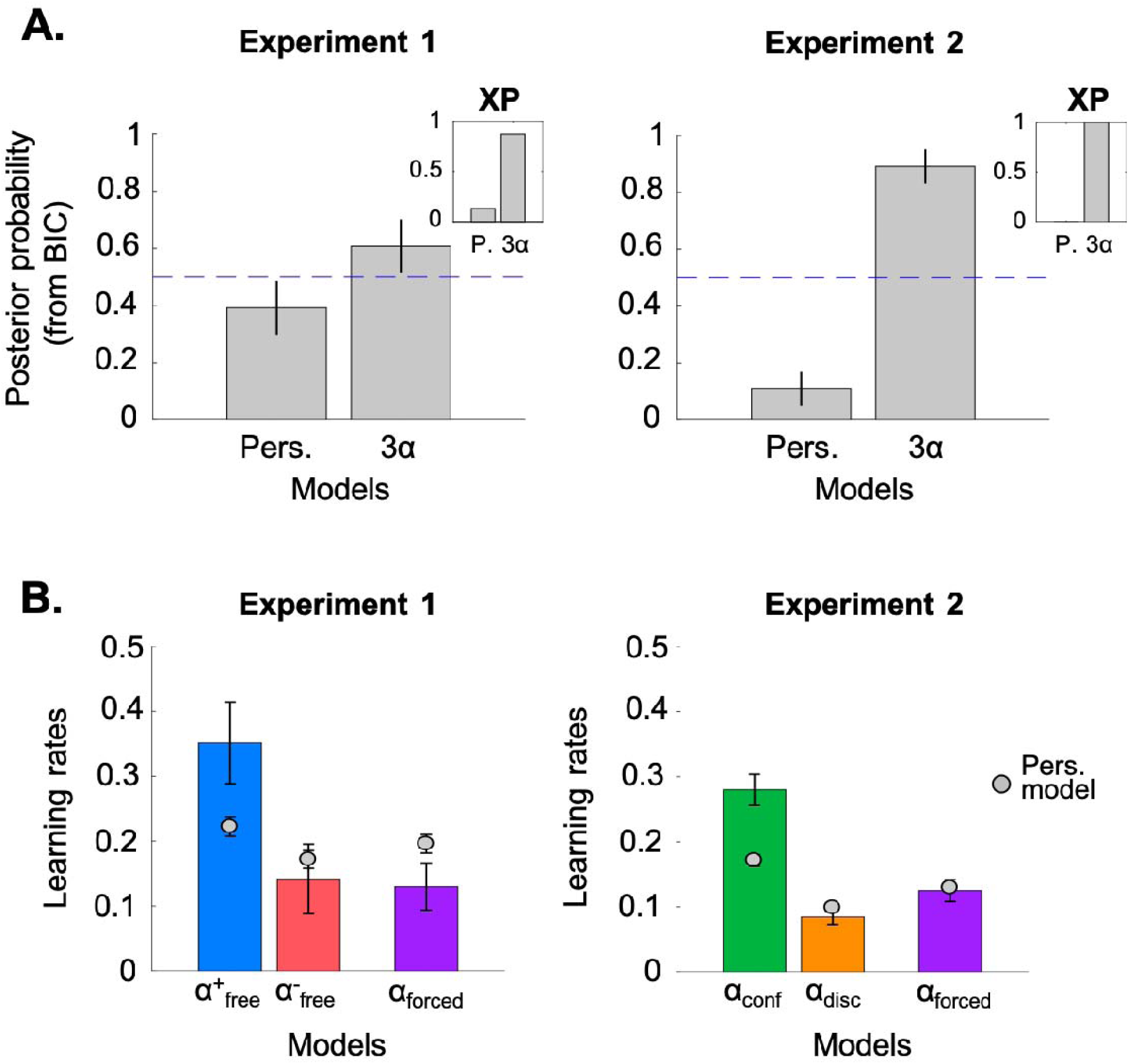
Comparing the winning model (3α) to a model with a simple perseveration parameter (“Pers.”). **(A)** Expectation and variance of the posterior probability for the perseveration model and the winning model with 3 learning rates, based on the Bayesian Information Criterion (BIC) values. The insert chart shows the exceedance probability (XP) for each model. **(B)** The winning model was fitted on data from Experiment 1 (left panel) and Experiment 2 (right panel). The bars represent the learning rates fitted on the participants’ data, while the grey dots represent the learning rates fitted to artificial datasets, created by simulating the preservation model 10 times on each participant’s data. As can be seen on the figure, positivity (left) and confirmation (right) biases may spuriously arise from data featuring a perseveration bias, but the biases retrieved from the simulations (grey dots) were significantly smaller compared to those observed in the participants’ data (bars).

To further quantify the extent to which the observed learning biases could be ascribed to the observed choice repetition bias, we simulated the perseveration model (using its best fitting parameters) and fitted the models with multiple learning rates on these synthetic data (**Figure 6B**). While model parameter analyses confirmed that positivity and confirmation biases may spuriously arise from data featuring a perseveration bias, the biases retrieved from the simulations were nonetheless significantly smaller compared to those observed in the participants’ data (Experiment 1: *t_23_* = 2.18; *p* = 0.03; Experiment 2: *t_23_* = 5.56, *p* < 0.001).

## Parameter adaptation to task contingency

In the first two experiments, we manipulated reward contingencies to include a ‘low-reward’ (reward probabilities set to 0.4 and 0.1) and a ‘high-reward’ condition (reward probabilities set to 0.9 and 0.6). This manipulation was included to first assess whether learning rates were adaptively modulated as a function of the reward contingencies, and second, to test whether this modulation extended to forced-choice outcomes. Previous optimality analyses suggest that a positivity bias would be advantageous in low-reward conditions, while the opposite would be true in high-reward conditions (Cazé and van der Meer, 2013). In other terms, it would be optimal to exhibit a higher learning rate for *rare* outcomes (i.e., rewards in ‘low-reward’ conditions and punishments in ‘high-reward’ conditions). In a new computational analysis, we fitted different learning rates for high- and low-reward conditions, thus creating new models with 6 learning rates (3 (learning rate types) × 2 (low, high)). We subjected the resulting parameter values to a 3 (learning rate types) × 2 (high-vs. low-reward conditions) repeated-measures ANOVA.

As expected, the learning rate type had a significant main effect, although it was only marginal in Experiment 1 (Exp. 1: *F_1,23_* = 3.6, *p* = 0.061; Exp. 2: *F_1,23_* = 8.8*, p* = 3.7 × 10^-3^) (**Figure 7A**). Importantly, there was no main effect of the condition factor in both experiments (Exp. 1: *F_1,23_* = 0.067, *p* = 0.80; Exp. 2: *F_1,23_* = 1.0*, p* = 0.31). Regarding the condition-by-type interaction, the effect was equivocal: it was marginally significant in Experiment 1, but null in Experiment 2 (Exp. 1: *F_1,23_* = 3.1, *p* = 0.082; Exp. 2: *F_1,23_* = 0.27*, p* = 0.60; see Figure 8A). To further support the absence of evidence in favour of learning rate adaptation we turned to model comparison. The models with different learning rates for high- and low-reward conditions were compared to models without learning rate modulation (“pooled” models). In both experiments, we found that the model *without* reward-probability dependent learning rates had the highest exceedance probability (XP=1.00) (see **Figure 7B**).

**Figure 7.**
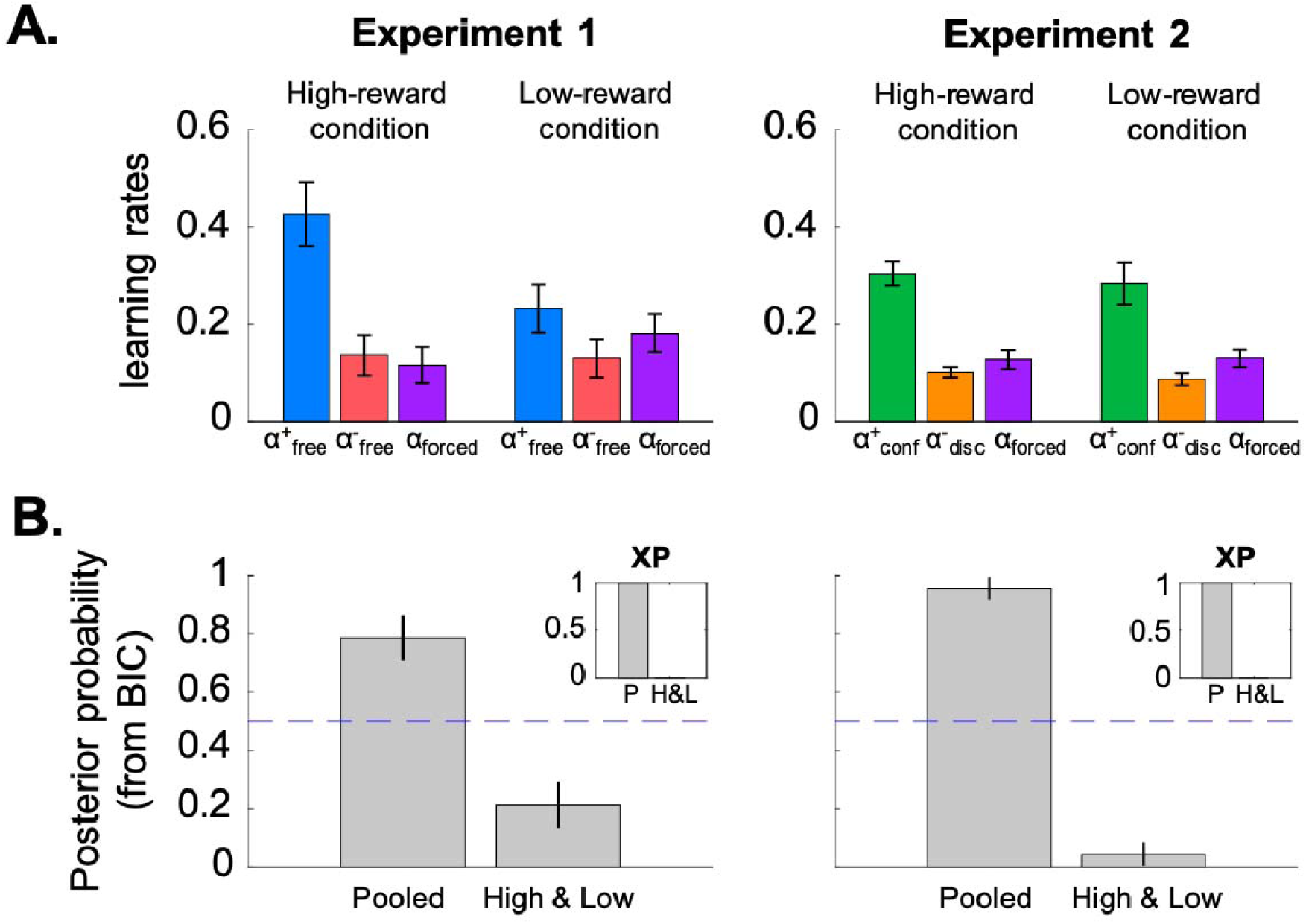
Learning rate analysis and model comparison for “High & Low” models. **(A)** The best-fitting learning rates of the “High & Low” model. **(B)** The posterior and exceedance probabilities of the “Pooled” (P) and “High & Low” (H&L) models. In contrast to the High & Low model, learning rates of the “Pooled” model are not modulated by reward contingencies.

**Figure 8.**
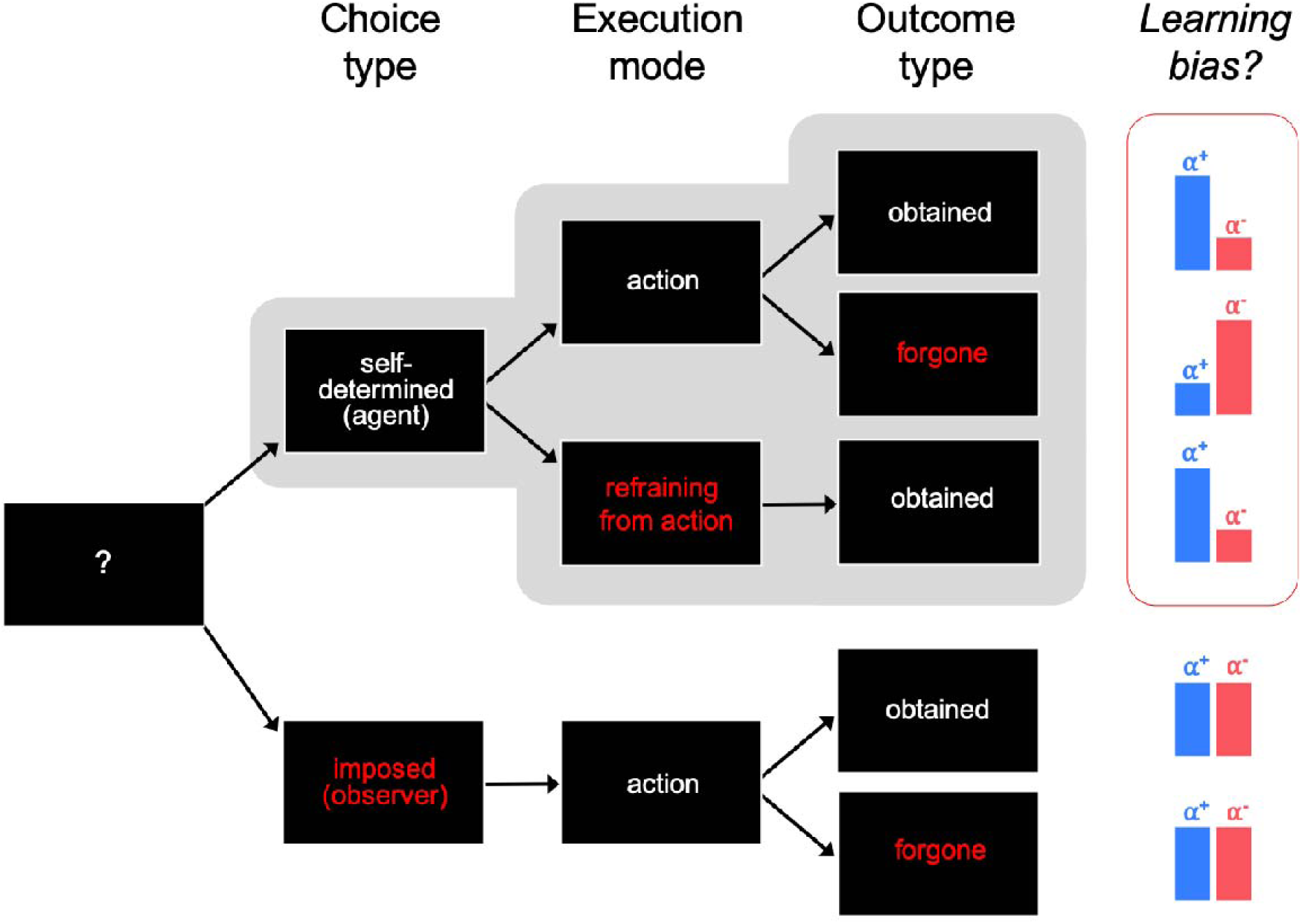
Valence-dependent learning biases as a function of the choice type, execution mode and outcome type. *Top: self-determined choice stream*. A valence-dependent learning asymmetry only arises when the individual is an agent, i.e., she “controls” the choice made. *Positive* (obtained) outcomes are better integrated than negative (obtained) outcomes – blue and red bars, respectively. This pattern reverses when learning from *forgone* outcomes, demonstrating that learning asymmetry reflects a *choice*-confirmation bias rather than a true positivity bias. Note this bias emerges early in the action processing chain, i.e., at the decision (or choice) rather than at the action stage. *Bottom: imposed choice stream*. No learning asymmetry is observed when the individual is not an agent, i.e., she does not control the choice made but only matches the computer’s decision. In this situation, participants learn from positive and negative outcomes alike.

## Discussion

While standard accounts of belief (or value) prescribe that an agent should learn equally well from positive and negative information (e.g., Barto and Sutton, 1998; Benjamin, 2018 for a review), previous studies have consistently shown that people exhibit valence-induced biases (e.g., Lefebvre et al., 2017; Kuzmanovic et al., 2018). These biases are traditionally exemplified in the so-called “good news/bad news” effect whereby people tend to overweight good news relative to bad news when updating self-relevant beliefs (Sharot and Garrett, 2016). An important moderator of these self-related biases would be the extent to which an individual believes she/he is able to *control* (i.e., choose) the dimension concerned. Thus, it has been shown that individuals tend to rate themselves as above average on positive *controllable*, but not uncontrollable, traits (Alicke and Govorun, 2005). Likewise, people would self-attribute positive outcomes when perceived controllability of the environment is high (Harris and Osman, 2012), and would enact different behaviours (Ajzen, 2002) or process differently behavioural consequences (Kool et al., 2013) when these consequences are under their direct control, relative to uncontrollable self-relevant outcomes (e.g., an asthma attack). In the present study, we sought to investigate further the link between outcome processing and control (i.e., voluntary choice) in three instrumental learning experiments comparing trials with and without voluntary choices (Exp. 1 & 2), featuring factual and counterfactual learning (Exp. 2), and implementing distinct action requirements (Exp. 3).

In the first experiment, learning performance was compared between trials in which the subject could either freely choose which option to select, or was “forced” to match a computer’s choice. As predicted, we found that participants learned better and faster from *positive*, relative to negative, prediction errors (PE), i.e., from better-than-expected outcomes. Crucially, this learning asymmetry (positive > negative PE) was present when participants were *free* to choose between options, but absent, if not reversed (negative > positive), when participants were *forced* to match an external choice. In other terms, we observed a positivity bias, i.e., a learning bias in favour of positively valued outcomes, only when learning was driven by *self-determined* choices.

In the second experiment, we combined free- and forced-choice trials with learning from factual (*chosen* action) or counterfactual (*unchosen* action) outcomes. Replicating previous results (Palminteri et al., 2017), we observed that prediction error valence biased factual and counterfactual learning in opposite directions: when learning from obtained outcomes (*chosen* action), positive prediction errors were preferentially taken into account than negative prediction errors. In contrast, when learning from forgone outcomes (*unchosen* action), participants integrated equally well positive and negative prediction errors. In other words, only *positive* outcomes that supported the participant’s current choice (positive outcomes associated with the *chosen* option) were preferentially taken into account, whereas positive outcomes that contradicted this choice (notably, positive outcomes associated with the *unchosen* option) were discounted. Taken together, these findings suggest that the well documented “positivity bias” may be a special case of a more general “choice-confirmation bias”, whereby individuals tend to overweight positive information when it confirms their previous choice. When no choice is involved, in contrast, positive and negative information are weighted equally (**Figure 8**).

Importantly, if learning asymmetry does reflect a choice-confirmation bias, then it should arise from the very first stage of the action processing chain, that is at the *decision* stage rather than at the *action* stage. Thus, a pure choice-supportive bias should be oblivious to how the choice is *implemented*, e.g., either through performing an action or refraining from acting. In contrast, a “cognitive dissonance” account would state that making an action is, in and of itself, sufficient to induce a learning asymmetry. Indeed, in the absence of any prior intention or reason to act, the mere fact of producing an action would strongly commit the agent with regard to the outcome. This commitment would then be retrospectively justified by shaping the individual’s preferences in such a way that they align *post hoc* with subsequent action outcomes (post-action dissonance reduction, see Izuma et al., 2010). Critically, most of the paradigms confound choice and action execution, and hence are not well suited to disentangle the influence of choice and action execution on valence-dependent learning asymmetries. In the third experiment, we directly addressed this issue by varying action requirements across “go” and “no-go” trials. Learning rates were analysed as function of both outcome valence (negative vs. positive) and execution mode (go vs. no-go). We replicated learning asymmetries found in free choice trials of both experiments 1 and 2, with positive PE being taken into account more than negative PE. Crucially, we found no difference between trials where the response was made by performing an action (key press) or by *refraining* from acting (no key press). Both dimensionality reduction and model comparison procedures supported these results. Thus, the choice confirmation bias is truly related to choices, rather than to the physical motor events that implement those choices.

Previous studies have suggested that learning rate asymmetries naturally implicate the development of choice *hysteresis*, where subjects tend to repeat previously rewarded options, despite current negative outcomes (e.g., Lefebvre et al., 2017). However, the very same choice behaviour may, in principle, derive from choice *inertia*, i.e., the tendency to repeat a previously enacted choice (Lau and Glimcher, 2005; Katahira, 2018). To settle this issue, we directly compared these two accounts and found that choice hysteresis was overall better explained by a choice-confirmation bias is terms of both model comparison and parameter retrieval. Thus, in our task we suggest that choice perseveration is better explained as biases in learning and updating. However, this does not exclude the possibility that learning-independent choice perseveration plays a role in decision-making, for instance, when the same pairs of cues are presented at the same spatial position across a higher number of trials.

Interestingly, theoretical simulations have suggested that preferentially learning from positive or negative prediction errors would be suboptimal in most circumstances, being only advantageous under specific and restrictive conditions – i.e., in environments with extremely distributed resources. Thus, Cazé and van der Meer (2013) demonstrated that – over the long run – different learning rate asymmetries can be advantageous for certain reward contingencies, which they referred to as “low-reward” and “higher-reward” conditions (i.e., the reward probabilities associated with the two available options are both low, or both high, respectively). Consistent with previous reports, we found no detectable sign of learning rate adaptation as function of reward contingencies (Palminteri 2017; Gershman, 2015). The absence of reward-dependent learning rates, if confirmed, is actually at odds with the above-mentioned optimality analysis, positing that it is more advantageous to have a lower positive than negative learning rate in high-reward conditions.

At first sight, it may be surprising that a “biased” model best accounts for participants’ data in a task where the rewards are dependent on participants’ performance. As a matter of fact, confirmation bias has been implicated in various suboptimal decisions, from wrongful convictions in judicial process (Findley and Scott, 2006) to academic underachievement (Rosenthal and Jacobson, 1968) and misinterpretation of scientific evidence (Loehle, 1987). While biased learning may be suboptimal locally or under specific conditions (e.g., being overly pessimistic about the consequences of other people’s decisions), it could be on average well suited to adapt to periodically changing environments. In real-world situations, both the amount of resources and the causes that bring about these resources – as when one is free to choose vs. forced to take actions under influence or coercion – may vary from time to time. Overweighting positive consequences resulting from voluntary choices, while keeping impartiality when acting under influence, might be the most robust pattern to deal with the intrinsic volatility of disposable resources (low/high) as well as the variety of their causal sources (internal/external). Accordingly, we found that our best-fitting model (choice-confirmatory in free choices, valence-neutral in forced-choices) was not only very advantageous in terms of accuracy, but it also exhibited the most stable performances across *low-* and *high-reward* conditions, relative to other models with alternative patterns of learning rates (see Supplementary Results, **Figure S2**).

Previous discussions of confirmation bias often focussed on “person-level” constructs, such as self-esteem, self-confidence, and post-decisional dissonance. However, we additionally suggest that a choice-confirmation bias could be adaptive in the context of the natural environment in which the learning system evolved (Fawcett et al., 2014). Previous accounts have highlighted the numerous benefits and facilitative effects of self-determined (vs. forced) choices on learning performance (e.g., Murayama et al., 2013). Besides enhancing memory encoding and retention (e.g., Voss et al., 2011) and boosting selectivity to choice-consistent evidence (Talluri et al., 2018), making self-determined choices improves learning of perceptual categories (e.g., Markant and Gureckis, 2010) and would allow for better generalization of prior knowledge to novel objects and situations (e.g., Wu and Tenenbaum, 2007). Choice allows people to control the stream of evidence they experience, and hence to focus effort on information that aligns with their current needs or interests, resulting in better and better-targeted learning (Gureckis and Markant, 2012, for a review). Choice is a powerful instrument to manipulate the environment so as to satisfy an individual’s needs (Leotti et al., 2010). A choice-confirmation bias would lead to preferentially reinforce actions that are most likely to meet these needs, i.e., freely chosen actions. In contrast, outcomes obtained from “forced” actions should be treated impartially as they do not necessarily align with the individual’s needs, interests or values, and hence should not be assigned any special value in self-determined decisions (Cockburn et al., 2014).

Our results bear intriguing resemblance with recent findings on self-attribution in causal inference. In a reinforcement learning task manipulating the probability of hidden-agent interventions, Dorfman and colleagues showed that when a participant believes that a benevolent agent has intervened, she learns more from negative than positive outcomes (i.e., she infers that the negative outcome is a consequence of her own choice rather than due to the benevolent agent). Conversely, when she believes that an adversarial agent has intervened, the participant is more likely to learn from positive than negative outcomes (Dorfman et al., 2019). These findings highlight the relation between valence-induced learning biases and control beliefs, and support the notion that people interpret feedback/changes in the environment differently according to perceived controllability. Controllability is a possible auxiliary hypothesis for interpreting changes in the environment (Gershman, 2019). Thus, if controllability is high, then negative outcomes are presumably not a consequence of one’s enacted choice, and so are underweighted (optimistic belief). In our task, likewise, we found that people overweight positive outcomes when selection of an option was under their direct control (free choice), yet are more impartial when they simply implement an instructed choice. Note that both Dorfman’s findings and ours are consistent with the notion that optimistic bias does not exclusively reflect preference for positive events in general – hence is not only a consequence of increased salience of positive outcomes. Rather, it would reflect biased (control) beliefs about one’s own causal power and the controllability of the environment (see also Chambon et al., 2018). Different beliefs about controllability might account for commonly observed differences between internal and external attribution profiles (Rotter, 1954) as well as between optimistic and pessimistic explanatory styles (Abramson et al., 1978).

## Conclusions

In three studies mixing free- and forced-choice trials, featuring both factual and counterfactual learning, and implementing distinct action requirements, we showed that participants’ behaviour was best accounted for by a learning model featuring a *choice-confirmation* bias – i.e., a model amplifying positive prediction errors in free-choice trials while being valence-neutral on forced-choice trials. We suggest that such a bias could be adaptive in the context of the natural environment in which the learning system evolved. Voluntary choices allow individuals to focus effort on information that aligns with their current needs. A choice-confirmation bias would thus lead to preferentially reinforce *freely chosen* actions, which are most likely to meet these needs. In contrast, outcomes obtained from *unchosen* (“forced”) actions should be treated impartially as they do not necessarily align with the individual’s needs, interests or values, and hence should not be assigned any special value in self-determined decisions. Interestingly, choice can be seen as an opportunity to exert control over the environment. Our results support the notion that people interpret action feedback differently depending on how controllable their environment is, in line with previous findings about self-serving bias in causal inference.

## Methods

### Participants

The study included 3 experiments. Each experiment involved 24 participants (Experiment 1: 13 males, mean age = 25.1 ± 0.8; Experiment 2: 9 males, mean age = 23.9 ± 0.5; Experiment 3: 10 males, mean age = 24.8 ± 0.7, **Table 1**). The local ethics committee approved the study (CPP C07-28). All participants gave written informed consent before inclusion in the study, which was carried out in accordance with the declaration of Helsinki (1964, revised 2013). The inclusion criteria were being older than 18 years, reporting no history of neurological or psychiatric disorders and a normal or corrected-to-normal vision. Participants were paid 10, 15 or 20 euros, depending on the number of points they had accumulated during the experiment.

### General procedure

Participants performed a probabilistic instrumental learning task modified from previous studies (Lefebvre et al., 2017; Palminteri et al., 2017). The task required choosing between two symbols that were associated with stationary outcome probabilities. The possible outcomes were either winning or losing a point. Participants were encouraged to accumulate as many points as possible and were informed that one symbol would result in winning more often than the other. They were given no explicit information about the exact reward probabilities, which they had to learn through trial and error.

Participants were also informed that some trials (indicated by the word “observer” displayed in the centre of the screen) were purely observational: the observed outcome would not be added to the total of points obtained so far, but it would allow them to gain knowledge about what would have happened should they have chosen the selected symbol (**Figure 1A**). Crucially, in forced-choice trials, the two symbols were pseudo-randomly preselected, thus ensuring equal sampling from both low- and high-reward options. In Experiment 2, participants were also informed that in some blocks, they would see the outcome associated with the unchosen symbol, although they would only accumulate the points associated with the chosen symbol (**Figure 1A**). In Experiment 3, the “observer” manipulation was not included, but subjects were instructed to express their decision by either making a key press (“go” trials) or refraining from making a key press (“no-go” trials) (**Figure 1B**).

### Conditions

Experiment 1 and 2 included four types of trials. In “free-choice” trials, participants could freely select between two possible symbols, while in “forced-choice” trials, participants had to match a preselected option. In “partial” trials, participants were only shown the outcome (“+1” or “-1”) associated to the chosen option, while in “complete” trials, participants were shown the outcome of *both* the chosen and unchosen symbols (**Figure 1**). The Experiment 1 included two conditions: a condition with only “partial” free-choice trials (40 trials per block), and a condition with intermixed “partial” free- and forced-choice trials (40 + 40 = 80 trials per block). In this “intermixed” condition, the free- and forced-choice trials were pseudo-randomly presented within the block, and the same pair of symbols was used in both types of trial.

In addition to the condition with intermixed partial free- and forced-choice trials, the Experiment 2 also consisted in a condition with intermixed “complete” free- and forced-choice trials. For the sake of duration, the number of trials was halved in Experiment 2 (20 free-choice trials + 20 forced-choice trials = 40 trials per block) (see **Table 1**).

Experiment 3 consisted of two “free-choice” conditions: a “go” and a “no-go” condition in which half of the participants had to press a computer key to select the *top* symbol, and to refrain from pressing any key to select the *bottom* symbol (the converse for the other half: press=bottom, refrain=top).

In Experiment 1, participants underwent 12 blocks of either 40 (free) or 80 (intermixed free+forced) trials each. Six blocks were “high-reward” blocks and six blocks were “low-reward” blocks. In high-reward blocks, one of the two symbols was associated with a .9 probability of winning (+1 point) – and hence with a .1 probability of loss (-1 point). The other symbol was associated with a .6 probability of winning. In low-reward blocks, one symbol was associated with a .4 probability of winning and the other with a .1 probability of winning. In Experiment 2, each condition consisted in 8 blocks of 40 (intermixed free+forced) trials each. Half of them were high-reward blocks. The low- and high-reward blocks were associated with the same reward contingencies as in Experiment 1. In Experiment 3, participants underwent 6 blocks of 100 (intermixed Go+No-Go) trials each. Since the issue concerning the adaptive modulation of learning rates was addressed in Experiments 1 and 2, we switched back to the contingencies used in a previous study (Palminteri et al., 2017). Thus, in half of the blocks, one symbol was associated with .7 probability of winning (+1 point) – and hence with a .3 probability of loss (-1 point) – and the other symbol was associated with a .3 probability of winning. In the other half, each symbol was associated with .5 probability of winning and losing (i.e., there was no correct response in these blocks).

In all experiments, each block was associated with a specific pair of symbols, meaning that the participant had to learn from scratch the reward contingencies at the beginning of each block. The first block was preceded by a short training (60 trials for Experiment 1; 40 trials for Experiment 2; 40 trials for Experiment 3). To ensure participants would not be biased toward expecting more frequent positive or negative outcomes in the subsequent experiment, the reward probabilities were set to .5 for all symbols during the training block.

### Trial structure

In Experiments 1 and 2, trials began with a fixation cross, except when free- and forced-choice trials were intermixed, in which case the words “actor” or “observer” appeared for 1000ms prior to each trial, depending on the type of choice involved (free- or forced-choice, respectively) (**see Figure 1A**). A pair of symbols was then presented in the left and right part of the screen (pseudo-randomly assigned on each trial). Participants made their choice by pressing the right or left key arrow with their right hand.

In forced-choice trials, the preselected cue was surrounded by a square. Participants had to press the corresponding arrow in order to move to the subsequent trial (nothing happened if they tried to press the other arrow). The cues were preselected to ensure equal sampling: one symbol was preselected in half of the trials, and the other symbol in the remaining trials. The obtained outcome was then presented in the same part of the screen as the chosen symbol. In “complete” trials, the foregone outcome was shown in the same part as the unchosen symbol. In intermixed free- and forced-choice trials, to ensure that participants paid attention to the outcomes presented, they were asked to press the up key arrow when winning a point and the down key arrow when losing a point.

In Experiment 3, trials began with a fixation cross for 1000ms (**Figure 1B**). A pair of symbols was then presented at the top or bottom of the screen (pseudo-randomly assigned on each trial). Participants had 1500ms to press the instructed key: the up key arrow for half of the participants, and bottom key arrow for the other half (Go trials). If no key was pressed after that delay, the other symbol was automatically selected (No-Go trials). In both Go and No-Go trials, a feedback associated with the selected symbol was then displayed for 1500ms.

### Computational modelling

We fitted the data with modified versions of a Q-learning model, including different learning rates following positive and negative prediction errors, and different learning rates in free- and forced-choice trials, or in Go and No-Go trials (see below). For each pair of symbols, the model estimates the expected value (also called Q-value) of the two options (Sutton & Barto 1991). The Q-values were set to 0 at the beginning of each block, corresponding to the a priori expectation of an equal probability of winning or losing 1 point. After each trial *t*, the value of the chosen option in a given state (*s*) is updated based on the prediction error, which measures the discrepancy between actual outcome value and the expected outcome for the chosen symbol, i.e., the chosen (*c*) Q-value, as follows:

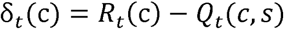

where *R_t_*(*c*) represents the obtained (factual) outcome on trial *t*.

The prediction error is then used to update the chosen Q-value:

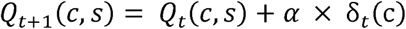

where α represents the learning rate parameter.

In the “complete” condition experienced in Experiment 2, participants could learn from both the obtained and the forgone outcomes. Thus, in these trials, the *unchosen* Q-value was also updated with the forgone (or unchosen: *u*) outcome using to the same rule:

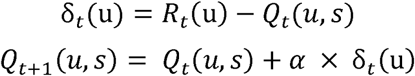

As mentioned above, different learning rates (α^+^ and α^-^) were fitted to reflect different updating processes after a positive or negative outcome (Lefebvre et al., 2017; Palminteri et al., 2017). *R* could take the following values *R* ϵ {−1, +1}, depending on the subject winning or not a point. The Q-values were initialized as such *Q*(1) = 0, which corresponds to unbiased prior expectations, and to the average outcome experienced during the training phase. Because we were interested in the specific effect of choice type on learning, different pairs of asymmetrical learning rates in free- and forced-choice trials (Experiments 1 and 2), and for factual and counterfactual outcomes (Experiment 2 only), were also fitted. The “Full” model thus had 4 learning rates in Experiment 1, and 8 learning rates in Experiment 2. In Experiment 3, different pairs of learning rates for Go and No-Go trials were fitted. The full model therefore included 4 learning rates (as in Experiment 1).

In the reinforcement learning framework, the stimuli with the highest Q-value is more likely to be selected. The probability of selecting the stimulus with the highest value was estimated with a softmax function, as follows:

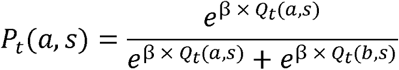

where β is the exploitation intensity parameter, which represents the strength of the Q-values on choice selection, and *a* and *b* being the two options available in a given state *s*. We fitted a unique parameter β for all trials and outcome types, to avoid biasing the learning rate comparison procedure. We also designed simpler versions of the full models in order to assess, for each experiment, what was the maximum number of parameters authorized, when penalizing for their complexity (parsimony-driven dimensionality reduction). The model space ranged from “full” models assuming different learning rates all possible outcomes (obtained and forgone), choice (free and forced) and action (go and no-go) types, to fully “reduced” model, assuming no bias at all.

### Perseveration model

Following Katahira’s critique (2017), suggesting that learning rates asymmetries may be artifactually be driven by a repetition (or perseveration) bias, we compared our models to a model including a ‘stickiness’ parameter. In the latter, the action selection rule was modified as follows:

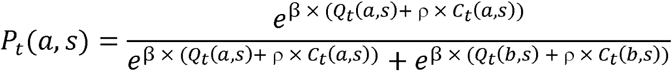

where the parameter ρ represents the participant’s tendency to perseverate, and *C_t_*(*x*, *s*) indicates which stimulus was chosen on the previous trial:

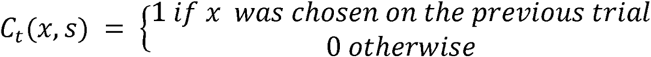

When the participants were forced to match the preselected option, we considered that they would not tend to perseverate in that choice. In the subsequent free trials, we thus set *C_t_*(*x*, *s*) to zero for both the preselected and the other stimuli.

### Parameter estimation

We fitted the model parameters based on participants’ choices on each free-choice trial, for each participant. We used a maximum posterior approach (or MAP, Bishop 2006) to avoid degenerate parameter estimates. The best parameters were those maximizing the logarithm of the posterior probability (LPP):

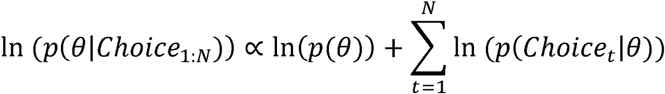

where θ represents our parameter set, *N* is the total number of trials in the experiment, and *p*(*Choice_t_*|*θ*) is the probability that the model would choose the same stimulus as the participant on trial *t*. To maximize the LPP with respect to θ, we used the Matlab’s “fmincon” function with the following ranges: 0 < β < Infinite, and 0 < α_i_ < 1.

The parameter prior probabilities were based on Daw et al. (2011), and we used a gamma distribution with hyperparameters 1.2 and 5 for the β parameter, and a beta distribution with hyperparameters 1.1 and 1.1 for the learning rates (α) parameters. To avoid biasing the learning rate comparison procedure, the same priors were used for all learning rates.

### Parameter recovery

We performed a parameter recovery analysis to ensure that the values of the learning rates reflected true differences in learning, and were not an artifact of the parameter fitting procedure (Palminteri et al., 2017). The aim was to check the capacity of recovering the correct parameters using simulated datasets. To do so, we first simulated performance on the two behavioral experiments using virtual participants. For each of these virtual participants, a learning rate value was being randomly drawn from a uniform distribution between 0 and 1. We then averaged the correlation coefficients *R* and *p*-values from 100 correlations performed between the parameters manipulated and the parameters recovered from the fitting procedure applied to the simulated data set (see Meyniel et al., 2016). This analysis was conducted on all the learning rate parameters of the full models (see **Supplementary Results, Figure S1**).

### Model comparison

The logarithm of the parameter posterior probability was used to compute the Bayesian Information Criterion (BIC; Schwarz, 1978) for each model and each participant, as follows:

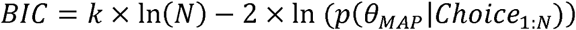

where *k* is the number of parameters, and (*p*(*θ_MAP_*|*Choice_1:N_*) is the logarithm of the posterior probability (LPP) of the MAP parameters given the participant’s choice data.

BIC of the different models were compared to verify that the extra learning rate parameters were justified by the data. As an approximation of the model evidence, individual BICs were fed into the MBB-VB toolbox (Daunizeau et al., 2014), a procedure that estimates how likely it is that a specific model generates the data of a randomly chosen subject (the posterior probability of a model) as well as the exceedance probability (XP) of one model being more likely than any other models in the set.

## Acknowledgements

V.C. was supported by the Agence Nationale de la Recherche (ANR) grants ANR-17-EURE-0017 (Frontiers in Cognition), ANR-10-IDEX-0001-02 PSL (program “Investissements d**’**Avenir”), and ANR-16-CE37-0012-01. H.T. was supported by a PSL/ENS studentship. P.H. was supported by the Chaire Blaise Pascal of the Région Île-de-France. S.P. was supported by a Marie Sklodowska-Curie Individual European Fellowship (PIEF-GA-2012 Grant 328822) and by an ATIP-Avenir grant.

## Author Contributions

S.P., V.C. and P.H. developed the study concept. Testing and data collection were performed by H.T. and M.V.. H.V. helped writing the PsychToolBox script for data collection. Data analysis was performed by H.T., M.V., V.C. and S.P.. V.C. and H.T. drafted the manuscript, and S.P. and P.H. provided critical revisions. All authors approved the final version of the manuscript for submission.

## Supplementary Material

### Parameter recovery

As a quality check for comparison between learning rates, we assessed parameter recoverability in both experiments 1 and 2 (see **Methods**, “Parameter recovery”). To do so, we fitted simulated datasets with known parameter values and found that learning rates were satisfactorily recovered on average (correlations on the diagonal: *R’s* > 0.78, all *p’s* < 10^-3^, **Figure S1**). Crucially, our fitting procedure introduced no spurious correlations between the true parameters and the recovered parameters (correlations outside the diagonal: -0.058 < *R’s* < 0.082, all *p’s* > 0.43, **Figure S1**).

**Figure S1.**
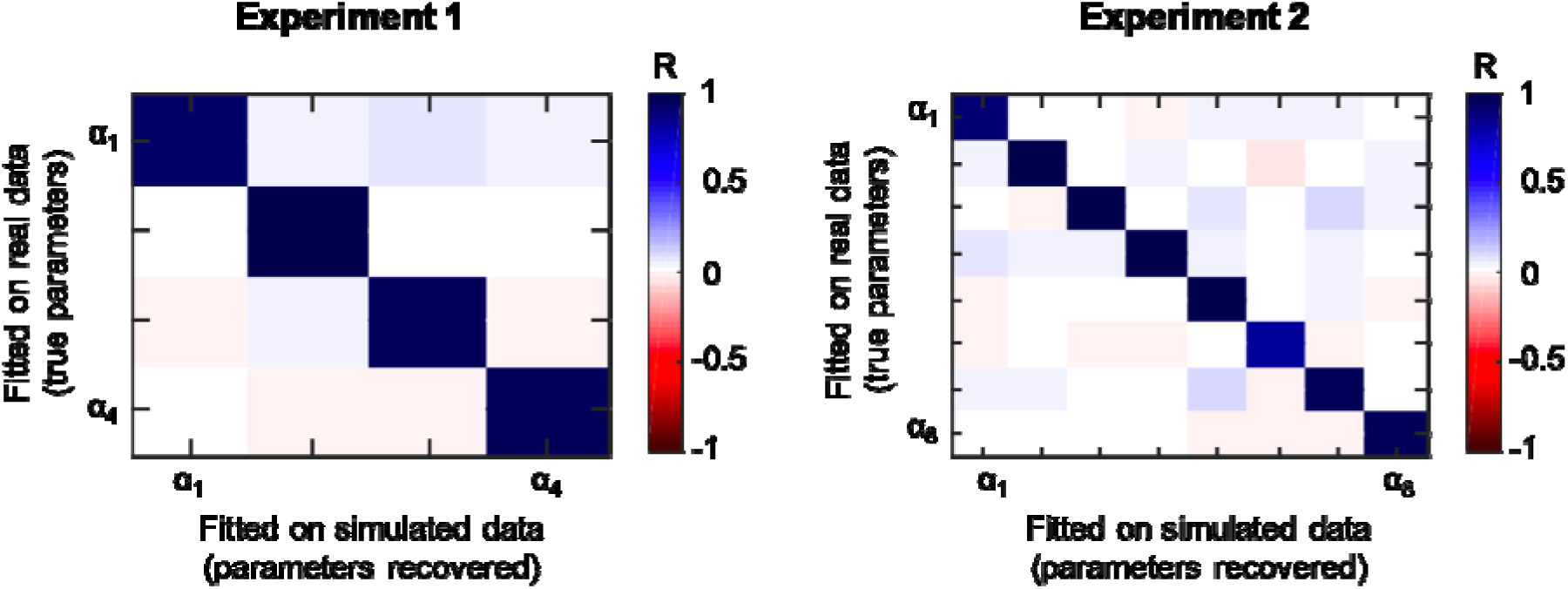
Parameter recovery results of the “full” models with 4 (Experiments 1) or 8 (Experiment 2) learning rates. The matrices show the averaged correlation coefficients (**R**). The correlations were computed between the parameters used to simulate artificial datasets (**y**-axis) and the parameters recovered from using our fitting procedure on these simulated datasets (**x**-axis).

### Model simulations

Using extensive model simulations, we assessed how the observed pattern of learning rates compares to other relevant patterns with respect to task performance (mean accuracy and variance).

We simulated models with different learning rate patterns on 1000 datasets for each participant to understand how parameter values could affect performance in our task. We set learning rates to be either choice-confirmatory (CO), or valence-neutral (NT) or choice-disconfirmatory (DC), and the learning rate patterns could be different in free-choice and forced-choice trials (see **Figure S2** and **Table S1**). Replicating Cazé and van der Meer’s findings (2013), we also found the choice-confirmatory (CO) models to outperform the other models in low-reward conditions, and the choice-disconfirmatory models to have better performances in the high-reward conditions (**Figure S2**).

When we looked at the general performance across both conditions, we found that the model corresponding to the participants’ learning rate patterns, i.e., the CO & NT model whose learning rates were choice-confirmatory in free choices, and valence-neutral in forced choices, was among the highest performing model (**Figure S2**). In Experiment 1, the CO & NT model had a performance of 83.2%, while the CO & DC model had a slightly higher performance of 84.2%. In Experiment 2, the CO & NT model had a performance of 86.6% while the CO & DC model had a slightly lower performance of 85.7%. Therefore, the learning rate patterns found in our participants can be said to be optimal, or close to optimal.

Interestingly, the performances of the CO & NT model were also quite similar across the high- and low-reward conditions. In Experiment 1, the difference in performances between high- and low-reward conditions was 2% for the CO & NT model and 1.9% for the CO & DC model, while this difference was over 3% for the other models. In Experiment 2, the difference in performances was the smallest for the CO & NT model (0.4% versus 0.8% for the CO & DC model, and 0.5% for the NT & CO model). Not only the participants’ best-fitting pattern was very advantageous in terms of accuracy, but it also exhibited highly stable performances across low- and high-reward conditions. Crucially, our participant also showed similar performances across both conditions (paired t-tests for Exp. 1: *t_23_* = 0.25, *p* = 0.80; for Exp. 2: *t_23_* = -0.027, *p* = 0.98).

**Figure S2.**
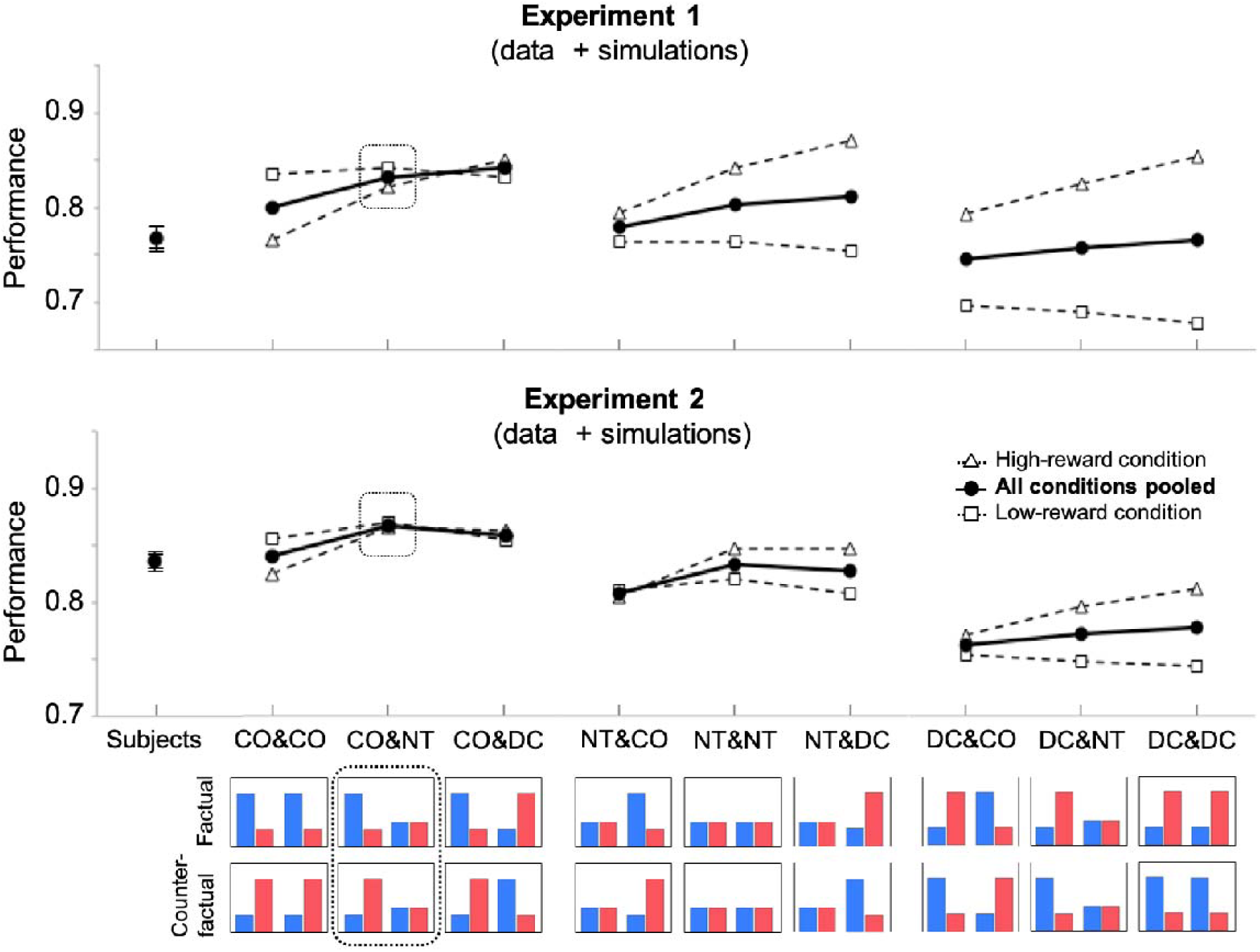
Parameter optimality was tested by simulating models with different learning rate patterns in Experiments 1 and 2. The models were labelled according to their learning rate patterns, shown in the bottom panel. For example, “CO & NT” designates a model with choice-confirmatory learning rates in free-choice trials, and valence-neutral learning rates in forced-choice trials. The diamonds and the squares correspond to the performance in high- and low-reward conditions, respectively. The black circles correspond to the performance averaged across the two conditions. Error-bars were plotted, although they were often too small to be seen. Bottom: the participants’ pattern of learning rates was highlighted with a dashed square.

**Table S1.** Learning rates used to simulate models with a choice-confirmatory (**CO**), a valence-neutral (**NT**) or a choice-disconfirmatory (**DC**) pattern on free- or forced-choice trials in Experiment 2.

|  | *Factual* |  | *Counterfactual* |  |
| --- | --- | --- | --- | --- |
| | $\alpha^+$ | $\alpha^-$ | $\alpha^+$ | $\alpha^-$ |
| **CO** | 0.3 | 0.1 | 0.1 | 0.3 |
| **NT** | 0.15 | 0.15 | 0.15 | 0.15 |
| **DC** | 0.1 | 0.3 | 0.3 | 0.1 |

## References

Abramson, L. Y., Seligman, M. E., & Teasdale, J. D. (1978). Learned helplessness in humans: Critique and reformulation. Journal of abnormal psychology, 87(1), 49.

Aberg, K. C., Doell, K. C., & Schwartz, S. (2016). Linking individual learning styles to approach-avoidance motivational traits and computational aspects of reinforcement learning. PloS one, 11(11), e0166675.

Bellebaum, C., Jokisch, D., Gizewski, E. R., Forsting, M., & Daum, I. (2012). The neural coding of expected and unexpected monetary performance outcomes: Dissociations between active and observational learning. Behavioural brain research, 227(1), 241–251.

Benjamin, D. J. (2018). Errors in probabilistic reasoning and judgment biases (No. w25200). National Bureau of Economic Research.

Bishop, C. M. (2006). Pattern recognition and machine learning. Springer.

Boureau, Y. L., & Dayan, P. (2011). Opponency revisited: competition and cooperation between dopamine and serotonin. Neuropsychopharmacology, 36(1), 74.

Brehm, J. W. (1956). Postdecision changes in the desirability of alternatives. The Journal of Abnormal and Social Psychology, 52(3), 384.

Burke, C. J., Tobler, P. N., Baddeley, M., & Schultz, W. (2010). Neural mechanisms of observational learning. Proceedings of the National Academy of Sciences, 107(32), 14431–14436.

Burke, C. J., Baddeley, M., Tobler, P. N., & Schultz, W. (2016). Partial adaptation of obtained and observed value signals preserves information about gains and losses. Journal of Neuroscience, 36(39), 10016–10025.

Caspar, E. A., Christensen, J. F., Cleeremans, A., & Haggard, P. (2016). Coercion changes the sense of agency in the human brain. Current biology, 26(5), 585–592.

Cazé, R. D., & van der Meer, M. A. (2013). Adaptive properties of differential learning rates for positive and negative outcomes. Biological cybernetics, 107(6), 711–719.

Chambon, V., Thero, H., Findling, C., & Koechlin, E. (2018). Believing in one’s power: a counterfactual heuristic for goal-directed control. bioRxiv, 498675.

Cooper, J. C., Dunne, S., Furey, T., & O’Doherty, J. P. (2012). Human dorsal striatum encodes prediction errors during observational learning of instrumental actions. Journal of cognitive neuroscience, 24(1), 106–118.

Correa, C. M., Noorman, S., Jiang, J., Palminteri, S., Cohen, M. X., Lebreton, M., & van Gaal, S. (2018). How the level of reward awareness changes the computational and electrophysiological signatures of reinforcement learning. Journal of Neuroscience, 38(48), 10338–10348.

Daunizeau, J., Adam, V., & Rigoux, L. (2014). VBA: a probabilistic treatment of nonlinear models for neurobiological and behavioural data. PLoS Computational Biology, 10(1), e1003441.

Daw, N. D. (2011). Trial-by-trial data analysis using computational models. Decision making, affect, and learning: Attention and performance XXIII, 23, 3–38.

Daw, N. D., Gershman, S. J., Seymour, B., Dayan, P., & Dolan, R. J. (2011). Model-based influences on humans’ choices and striatal prediction errors. Neuron, 69(6), 1204–1215.

Dorfman, H. M., Bhui, R., Hughes, B. L., & Gershman, S. J. (2019). Causal Inference About Good and Bad Outcomes. Psychological science, 0956797619828724.

Gershman, S. J. (2015). Do learning rates adapt to the distribution of rewards?. Psychonomic Bulletin & Review, 22(5), 1320–1327.

Gershman, S. J. (2019). How to never be wrong. Psychonomic Bulletin & Review, 1–16.

Filevich, E., Vanneste, P., Brass, M., Fias, W., Haggard, P., & Kühn, S. (2013). Brain correlates of subjective freedom of choice. Consciousness and Cognition, 22(4), 1271–1284.

Frank, M. J., Moustafa, A. A., Haughey, H. M., Curran, T., & Hutchison, K. E. (2007). Genetic triple dissociation reveals multiple roles for dopamine in reinforcement learning. Proceedings of the National Academy of Sciences, 104(41), 16311–16316.

Izuma, K., Matsumoto, M., Murayama, K., Samejima, K., Sadato, N., & Matsumoto, K. (2010). Neural correlates of cognitive dissonance and choice-induced preference change. Proceedings of the National Academy of Sciences, 107(51), 22014–22019.

Katahira, K. (2018). The statistical structures of reinforcement learning with asymmetric value updates. Journal of Mathematical Psychology, 87, 31–45.

Kuzmanovic, B., & Rigoux, L. Optimistic belief updating deviates from Bayesian learning. 2016. Available at SSRN: http://ssrn.com/abstract_id, 2810063.

Krieghoff, V., Brass, M., Prinz, W., & Waszak, F. (2009). Dissociating what and when of intentional actions. Frontiers in Human Neuroscience, 3, 3.

Lau, H. C., Rogers, R. D., Haggard, P., & Passingham, R. E. (2004). Attention to intention. Science, 303(5661), 1208–1210.

Lefebvre, G., Lebreton, M., Meyniel, F., Bourgeois-Gironde, S., & Palminteri, S. (2017). Behavioural and neural characterization of optimistic reinforcement learning. Nature Human Behaviour, 1, 0067.

Leotti, L. A., & Delgado, M. R. (2011). The inherent reward of choice. Psychological science, 22(10), 1310–1318.

Mather, M., Shafir, E., & Johnson, M. K. (2000). Misremembrance of options past: Source monitoring and choice. Psychological Science, 11(2), 132–138.

Mather, M., & Johnson, M. K. (2000). Choice-supportive source monitoring: Do our decisions seem better to us as we age?. Psychology and aging, 15(4), 596.

Mather, M., Shafir, E., & Johnson, M. K. (2003). Remembering chosen and assigned options. Memory & Cognition, 31(3), 422–433.

Meyniel, F., Goodwin, G. M., Deakin, J. W., Klinge, C., MacFadyen, C., Milligan, H., … & Gaillard, R. (2016). A specific role for serotonin in overcoming effort cost. Elife, 5.

Monfardini, E., Gazzola, V., Boussaoud, D., Brovelli, A., Keysers, C., & Wicker, B. (2013). Vicarious neural processing of outcomes during observational learning. PloS one, 8(9), e73879.

Murayama, K., Matsumoto, M., Izuma, K., Sugiura, A., Ryan, R. M., Deci, E. L., & Matsumoto, K. (2013). How self-determined choice facilitates performance: A key role of the ventromedial prefrontal cortex. Cerebral Cortex, 25(5), 1241–1251.

Murty, V. P., DuBrow, S., & Davachi, L. (2015). The simple act of choosing influences declarative memory. Journal of Neuroscience, 35(16), 6255–6264.

Nickerson, R. S. (1998). Confirmation bias: A ubiquitous phenomenon in many guises. Review of general psychology, 2(2), 175–220.

Niv, Y., Edlund, J. A., Dayan, P., & O’Doherty, J. P. (2012). Neural prediction errors reveal a risk-sensitive reinforcement-learning process in the human brain. Journal of Neuroscience, 32(2), 551–562.

Palminteri, S., Khamassi, M., Joffily, M., & Coricelli, G. (2015). Contextual modulation of value signals in reward and punishment learning. Nature communications, 6.

Palminteri, S., Lefebvre, G., Kilford, E. J., & Blakemore, S. J. (2017). Confirmation bias in human reinforcement learning: Evidence from counterfactual feedback processing. PLoS computational biology, 13(8), e1005684.

Palminteri, S., Wyart, V., & Koechlin, E. (2017). The importance of falsification in computational cognitive modeling. Trends in cognitive sciences, 21(6), 425–433.

Rotter, J. B. (1954). Social learning and clinical psychology.

Sharot, T., Velasquez, C. M., & Dolan, R. J. (2010). Do decisions shape preference? Evidence from blind choice. Psychological science, 21(9), 1231–1235.

Sharot, T., & Garrett, N. (2016). Forming beliefs: why valence matters. Trends in cognitive sciences, 20(1), 25–33.

Sutton, R. S., & Barto, A. G. (1991). Introduction to reinforcement learning. MIT Press.

Schwarz, G. (1978). Estimating the dimension of a model. The annals of statistics, 6(2), 461–464.

van Eimeren, T., Wolbers, T., Münchau, A., Büchel, C., Weiller, C., & Siebner, H. R. (2006). Implementation of visuospatial cues in response selection. Neuroimage, 29(1), 286–294.

Voss, J. L., Gonsalves, B. D., Federmeier, K. D., Tranel, D., & Cohen, N. J. (2011a). Hippocampal brain-network coordination during volitional exploratory behavior enhances learning. Nature neuroscience, 14(1), 115–120.

Yeung, N., Holroyd, C. B., & Cohen, J. D. (2004). ERP correlates of feedback and reward processing in the presence and absence of response choice. Cerebral cortex, 15(5), 535–544.

Yu, R., & Zhou, X. (2006). Brain responses to outcomes of one’s own and other’s performance in a gambling task. Neuroreport, 17(16), 1747–1751.

